# A population intrinsic timer controls *Hox* gene expression and cell dispersion during progenitor addition to the body axis

**DOI:** 10.1101/2023.05.10.540133

**Authors:** Lara Busby, Guillermo Serrano Nájera, Ben Steventon

## Abstract

During embryonic development, the timing of events at the cellular level must be coordinated across multiple length scales to ensure the formation of a well-proportioned body plan. This is clear during somitogenesis, where the progenitors must be allocated to the axis over time whilst maintaining a progenitor population for continued elaboration of the body plan. However, the relative importance of intrinsic and extrinsic signals in timing progenitor addition at the single cell level is not yet understood. Heterochronic grafts from older to younger embryos have suggested a level of intrinsic timing whereby later staged cells contribute to more posterior portions of the axis. To determine the precise step at which cells are delayed, we performed single-cell transcriptomic analysis on heterochronic grafts of somite progenitors in the chicken embryo. This revealed a previously undescribed cell state within which heterochronic grafted cells are stalled, post-ingression through the primitive streak. The delayed exit of older cells from this state correlates with expression of posterior *Hox* genes. Using grafting and explant culture, we find that both *Hox* gene expression and the migratory capabilities of progenitor populations are intrinsically regulated at the population level. Therefore, we demonstrate that cell dispersion is controlled by a population intrinsic timer to control progenitor addition to the presomitic mesoderm.

## Introduction

Timing is a fundamental concept in developmental biology: development is inherently dynamic and occurs with reproducible timing for the embryos of a given species (Busby and Steventon, 2021; Duboule, 2022; Ebisuya and Briscoe, 2018; Raff, 2006; Rayon, 2023). Developmental tempo refers to the cell-intrinsic features that produce species-specific developmental speeds including rates of transcription, translation, and protein turnover (Matsuda et al., 2020; Rayon et al., 2020). However, the mechanisms that control the absolute timing of developmental events within a particular embryo remain poorly understood. In the context of an embryo, cells which have intrinsic features must regulate their dynamics in accordance with their surroundings to produce emergent properties. The relative balance of intrinsic and extrinsic cues in timing events in development is an important question that has been investigated in contexts including limb development and the vertebrate tailbud (Chinnaiya et al., 2014; Fulton et al., 2021; Pickering et al., 2018; Saiz-Lopez et al., 2015; Saiz-Lopez et al., 2017).

The vertebrate primary body axis comprises a set of highly conserved axial and paraxial components, namely the midline notochord, dorsal neural tube, and pairs of somites on either side of the midline. These structures are produced in a defined anterior to posterior sequence, by the controlled allocation of axial progenitor cells to the body axis over time. In the avian embryo, this occurs through the passage of cells through the blastopore-analogous structures of the primitive streak and node, which will go on to form a structure termed the tailbud, found at the caudal extremity of the embryo (Brown and Storey, 2000; Catala et al., 1996; Gray and Dale, 2010; Iimura et al., 2007; McGrew et al., 2008; Schoenwolf, 1977; Selleck and Stern, 1991; Yang et al., 2002). Thus, a population of progenitor cells is maintained within the embryo throughout the period of axis elongation. The control of behaviour of these cells is key for population dynamics to be compatible with production of the full length of the body axis. Crucially, the contribution of cells from progenitor populations to the axis occurs in a gradual manner, with cells streaming through the primitive streak or later from the tailbud over a period of several days (Hamburger and Hamilton, 1951). The mechanisms which underlie this gradual streaming behaviour are unknown, though a role for *Hox* genes in timing axis contribution of progenitor cells has been proposed (Deschamps and Duboule, 2017; Iimura and Pourquie, 2006). During development of the vertebrate posterior body, axial progenitor cells express *Hox* genes in a differential manner over time – a phenomenon termed temporal collinearity (‘the Hox Clock’) (Deschamps and Duboule, 2017; Izpisúa-Belmonte et al., 1991). Specifically, as development progresses, axial progenitor populations express progressively more 5’ *Hox* gene complements. Previous work has implicated *Hox* genes in the control of cell behaviour during posterior body development: the over-expression of various 5’ *Hox* genes in axial progenitor cells of the avian primitive streak is sufficient to delay their contribution to the body axis (Iimura and Pourquie, 2006). More 5’ *Hox* genes delay the timing of cell contribution to a greater extent than more 3’ *Hox* genes (Iimura and Pourquie, 2006).

Importantly, the relative role of cell-intrinsic and extrinsic cues in timing the decision of individual cells to enter the body axis is unknown. The delayed contribution of heterochronic grafts of tissue from older to younger embryos relative to stage-matched grafts (Iimura and Pourquie, 2006; Tam and Tan, 1992) suggests a degree of intrinsic control over axis contribution timing, but the extent to which this is true remains to be understood. In this paper, we exploit the rich toolkit provided by experimental embryology (Busby et al., 2022) in conjunction with next-generation sequencing methods and multiplexed staining to investigate the extent to which axial progenitor gene expression and behaviour are intrinsically or extrinsically regulated.

A progenitor population that produces the medial portion of somites has been described in the avian embryo, residing at the 90% region of the primitive streak (PS) (Iimura et al., 2007; Psychoyos and Stern, 1996). We chose to focus on this cell population, herein termed the medial somite progenitor (MSP) population, as a model for understanding the timing of cell contribution to the body axis. In particular, we chose to focus our analyses on HH4 (definitive primitive streak stage) and HH8 (4 somite-stage) embryo, for several reasons. Whilst the somite progenitor population is continuous across these stages, it is known that the first 4-5 somites of the avian axis are distinct from more posterior ones – they are formed simultaneously (Dias et al., 2014), lack the rostro-caudal sub-patterning present in more posterior somites (Rodrigues et al., 2006), do not develop ganglia (Lim et al., 1987), and do not give rise to segmented structures but instead to the occipital and sphenoid bones of the skull (Couly et al., 1993; Huang et al., 2000). Cells of the HH4 MSP region give rise to the occipital somites (Psychoyos and Stern, 1996) whereas at HH8 these somites have formed already (Hamburger and Hamilton, 1951), and so the progenitor region exclusively produces trunk and tail somites. It is therefore interesting to consider possible differences between the dynamics and behaviour of the progenitor region between HH8 and HH4, and whether these are maintained irrespective of the environment that the tissue is placed in. In normal development, the context in which the HH4 and HH8 MSP regions allocate cells to the axis is very different – at HH4 (primary gastrulation) the mesendoderm represents an unbounded compartment that cells move into after ingression through the primitive streak whereas at HH8 a clear pre-somitic mesoderm is formed and cells are allocated to this bounded compartment. Despite the substantial differences between occipital and trunk somite formation, it remains unknown whether these modes of somitogenesis result from changing environmental inputs or changes to the progenitor population itself over time.

We find that heterochronic grafts of somitic progenitors from HH8 to HH4 are delayed relative to stage-matched grafts at a previously undescribed step that we term ‘dispersion’, occurring after cells pass through the primitive streak and enabling substantial mixing between donor and host tissue in the mesendodermal compartment. By culturing MSP tissue *ex vivo* on fibronectin, we find that the delayed migration is an intrinsic property of HH8 tissue and can account for the delay in axis contribution of heterochronic grafts. Surveying *Hox* gene expression in the MSP region at HH4 and HH8 reveals key differences in gene expression, which are maintained in grafted HH8 tissue in the HH4 environment, suggesting that *Hox* gene expression is also an intrinsic property of the tissue. Together, this work demonstrates that a population intrinsic timer controls Hox gene expression and cell dispersion during progenitor addition to the body axis.

## Results

### Delayed contribution of heterochronic grafts is characterised by a lack of dispersion of cells from the primitive streak

To ask whether the behaviour of progenitor cells differs before and after the onset of node regression (cessation of primary gastrulation), we compared homochronic HH4 to HH4 grafts with HH8 to HH4 grafts. Homochronic grafts of MSP cells were performed at HH4 (Hamburger and Hamilton, 1951) from a GFP transgenic line (McGrew et al., 2008) to a wild-type embryo (*Figure 1A*). Over time, the gradual contribution of cells deriving from the MSP population to the body axis can be observed (*Figure 1B-G*). At each timepoint, a population of MSP cells remains closely associated with the morphological remnant of the node during node regression after HH5 (indicated by the yellow boxes in *Figure 1B-G*), whilst other cells ingress through the streak and begin to migrate toward their final location in the medial portion of a somite (*Figure 1G’, G’’’*). At the final timepoint imaged (24 hours after grafting), GFP-expressing donor cells can still be found in the progenitor region, by which time the embryo has reached tailbud stage (*Figure 1G’’*).

**Figure 1:**
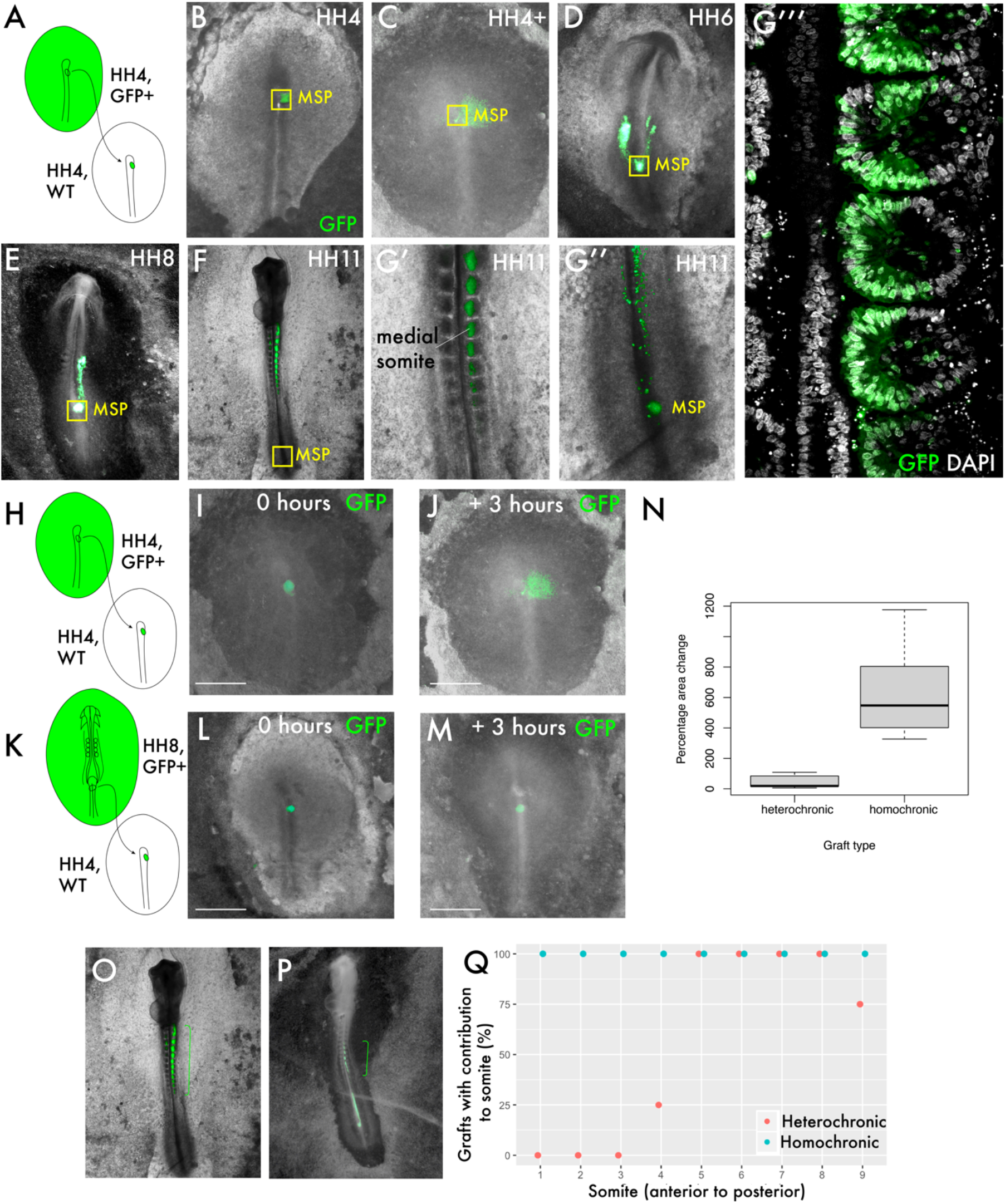
Delayed contribution of heterochronic grafts is characterised by a lack of dispersion of cells from the primitive streak. (A) Homochronic grafts of the 90% primitive streak (PS) region were performed from GFP-expressing HH4 embryos to wild-type HH4 embryos. (B)-(G’’) Combined GFP/brightfield images of grafts at various times between 0 and 24 hours after grafting reveal the behaviours of this population during body axis development. As the node regresses posteriorly after HH5, a subpopulation of the MSP region remains associated closely with the morphological remnant of the node (yellow box “MSP”), whilst other cells clearly part from this population and move toward the anterior of the embryo, where the somites will form. At 24 hours post-graft, embryos are approximately HH10-11 and GFP cells are clearly located in the medial portion of the somites (G’), close to the midline. At this timepoint, a population of GFP+ cells remain at the posterior progenitor region of the embryo (the tailbud) (G’’). (G’’’) is a single slice confocal image of the graft in (G’, G’’) stained with DAPI showing the precise localisation of grafted cells (GFP) in the medial portion of the somite. (H) and (K) are schematic representations of the two graft types, showing homochronic HH4-HH4 and heterochronic HH8-HH4 grafts respectively. (I) and (L) show overlaid GFP/ brightfield images of homochronic (HH4-HH4) and heterochronic (HH8-HH4) grafts at 0 hours after grafting. (I) and (L) show the same grafts at 3 hours, when the homochronic graft has spread substantially from the initial graft site but the heterochronic graft appears very similar to the 0-hour image. (N) Quantification of the change in area of GFP-positive grafted tissue in homochronic and heterochronic grafts at 3 hours vs. 0 hours post-graft, represented as a boxplot. (O) and (P) are combined GFP/ brightfield images of homochronic and heterochronic grafts, respectively, at 24 hours after grafting. The green brackets in (O) and (P) indicate the somite contribution of the two grafts. (Q) is a scatterplot showing the somite contribution of HH4-HH4 and HH8-HH4 grafts, with the percentage of grafts that have cells in each somite plotted against somite number. The different colours of datapoint represent the type of graft (homochronic or heterochronic). *Scale bars in (I), (J), (L) and (M) represent 1mm*.

We then performed grafts of the MSP region from HH8 embryos into the HH4 embryo (*Figure 1K*). Within 3 hours, GFP donor cells in homochronic grafts have spread appreciably from the graft site (*Figure 1I vs. 1J*), whereas the location of GFP donor cells in heterochronic grafts does not differ substantially from immediately post-grafting (compare *Figure 1L* with *Figure 1M*). The change in area occupied by the graft (between 0- and 3-hours post-grafting) was quantified and found to be statistically significant between the two graft types (*Figures 1N*). Importantly, this delay in dispersion correlates with a difference in anteroposterior somite contribution: whilst homochronic grafts invariably have their most anterior axis contribution in somite 1, heterochronic grafts typically have their most anterior contribution in somite 4 or 5 (*Figure 1O* vs. *1P*, quantification in *1Q*).

These results show that HH8 progenitor cells exhibit substantially different behaviour in the HH4 environment relative to stage-matched grafts as early as 3 hours after grafting. We therefore focused on the initial ingression and spreading of cells from the primitive streak to elucidate which cell state transitions are key for the timing of somite progenitor addition.

### Single cell analysis reveals a previously undescribed cell state within which heterochronic grafted cells are stalled

To investigate the cell states occupied by homochronic and heterochronic MSP grafts we turned to single cell RNA-sequencing (*Figure 2)*. Grafted tissue was dissected, pooled, and dissociated, with each sample comprising a mixture of GFP+ donor cells and GFP-host cells (*Figure 2A*). Cells from all three samples were integrated to a single dataset and clustered, yielding 14 distinct clusters (*Figure 2B*). Published expression data (GEISHA – Bell et al, 2004; Darnell et al., 2007) was used to annotate these cluster identities (*Supplementary Figure 1, Figure 2B, C*), and the expected cell types were present including epiblast, ingressed mesoderm, endoderm, and Hensen’s node. Note that there were four clusters in the dataset – 6, 8, 12 and 13 – which could not be annotated based on published gene expression. These clusters are found at the centre of the UMAP plot (*Figure 2B*). Splitting the dimensionality plot by sample (*Figure 2D-F*) shows some differences in the clusters present in each case – most notably, the HH4-4 sample lacks clusters 0, 4 and 6 which represent HH8 tissue (expressing posterior marker genes including *Hoxb8* and *Cdx4, Figure 2C*) (Barak et al., 2012; Bell et al, 2004; Darnell et al., 2007; Joshi et al., 2019). As the dataset comprises both host and donor cells in each instance, we plotted the expression of *eGFP* in each dataset to ask what is the distribution of donor tissue (*eGFP*+) compared to host cells (*Figure 2G-I*). In HH4-4 control grafts, *eGFP*-expressing cells are found across the clusters present in the dataset mixed with *eGFP*-negative wild-type cells, consistent with host and donor tissue being essentially equivalent in this instance (*Figure 2G*). In HH8-4 grafts at 0 hours (*Figure 2H*), *eGFP*-expressing cells are present in the left-hand clusters on the UMAP plot, clusters 0, 4, and 6. This shows that HH8 tissue occupies a distinct region in gene expression space initially upon grafting. Strikingly, after 3 hours, HH8 *eGFP*-positive cells are enriched in the central clusters 6, 8 and 12 (*Figure 2I*). Importantly, surveying the HH4-HH4 homochronic graft dataset shows that this cell state is not unique to the heterochronic graft condition but instead present in normal development, though there is not a particular enrichment of cells in this state (*Figure 2D, G*), suggesting that cells transit rapidly through this gene expression state during normal development. Thus, our scRNA-seq experiment reveals an undescribed cell state in mesodermal progenitor contribution to the body axis which HH8 cells accumulate in upon grafting.

**Figure 2:**
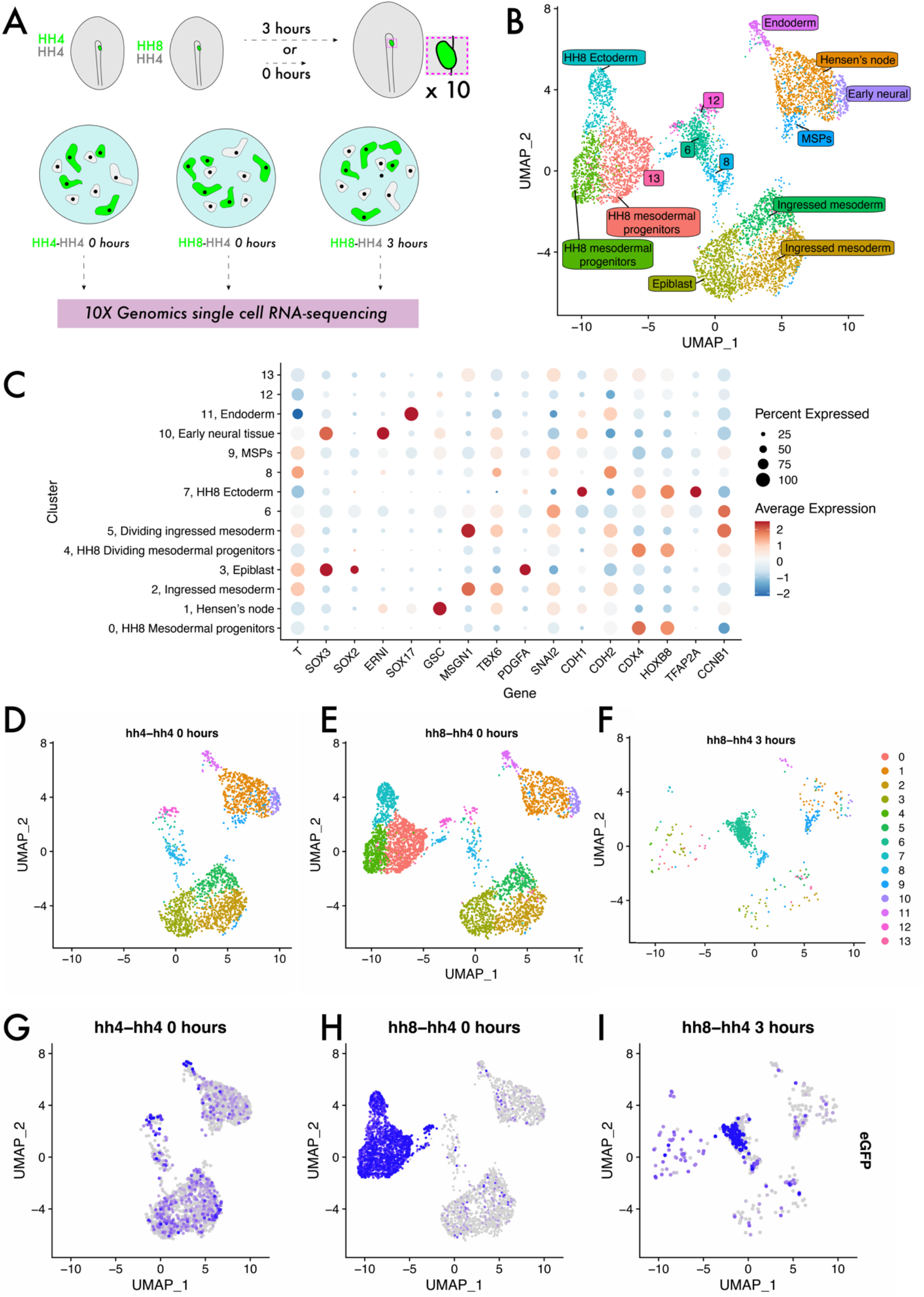
Single cell analysis reveals a previously undescribed cell state within which heterochronic grafted cells are stalled. (A) Schematic representing single cell RNA-sequencing experimental design. Grafts were performed from a GFP-expressing donor embryo to a wild-type host embryo and the region containing fluorescent tissue was dissected either 0 or 3 hours after grafting. 10 grafts were pooled for each condition and subjected to dissociation and single cell RNA-sequencing. (B) UMAP plot showing cells from all three datasets clustered according to gene expression differences. The colour of each point on the plot represents the cluster to which it has been assigned, and labels denote cluster identities assigned according to the expression of previously described ‘marker’ genes. Note that clusters 6, 8, 12 and 13 were unable to be assigned a specific cell state identity based on described gene expression. (C) Dot Plot showing the expression of genes in each cluster of the combined dataset. The appearance of each dot is a composite of the percentage of cells in that cluster with positive expression of the gene (size of dot) and the average quantitative level of gene expression (colour of dot) – see key on the far right of the plot. (D)-(F) UMAP plots of each sample with the colour of each spot (cell) representing the cluster to which it has been assigned. (G)-(I) UMAP plot of each sample with the colour of each spot (cell) representing the expression level of *eGFP;* more purple spots represent higher expression values. Cells with non-zero values can be identified as donor cells deriving from the GFP-expressing transgenic embryo.

### The transition from ingressed mesoderm to individual dispersed cells is a key step in gastrulation, associated with changes in cell adhesion

To ask what the cell states encompassed by clusters 6, 8 and 12 represent, we revisited our grafts to determine if heterochronically grafted cells had undergone an epithelial-to-mesenchymal transition (EMT) and ingressed through the primitive streak, or whether they remained in the epiblast. Re-slicing Z-stacks of these grafts allowed us to survey the localisation of eGFP-positive cells in the dorsoventral axis (*Figure 3A, B*). In both homochronic and heterochronic grafts, eGFP-positive cells clearly reside in the lower, ventral mesendodermal cell layer rather than the more dorsal epiblast (*Figure 3A, B*). When we surveyed the expression of *Snail2 (Snai2*) - considered a master regulator of EMT and required for mesodermal cells to ingress through the primitive streak (Nieto et al., 1994) - we found its expression in cells of clusters 6, 8 and 12 (*Figure 3D*, compare with *3C*). Indeed, we confirmed the expression of the EMT-associated transcripts *N-Cadherin (Cdh2*) (Dady et al., 2012) and *Snai2* in grafted tissue by multiplexed HCR (*Figure 3E, F*) and found their expression in both homochronic and heterochronic grafts at 3 hours. Taking the results in *Figure 1* with this data, this leads us to the conclusion that HH8 cells are delayed after ingression through the primitive streak at the level of a step we term *cell dispersion* – the spreading of cells in the AP and ML axes of the mesendoderm post-ingression (schematic in *Figure 3G*). Rather than mixing extensively with host tissue as in HH4-HH4 grafts, HH8 cells remain as a single group after grafting (compare *Figure 1J* and *IM*).

**Figure 3:**
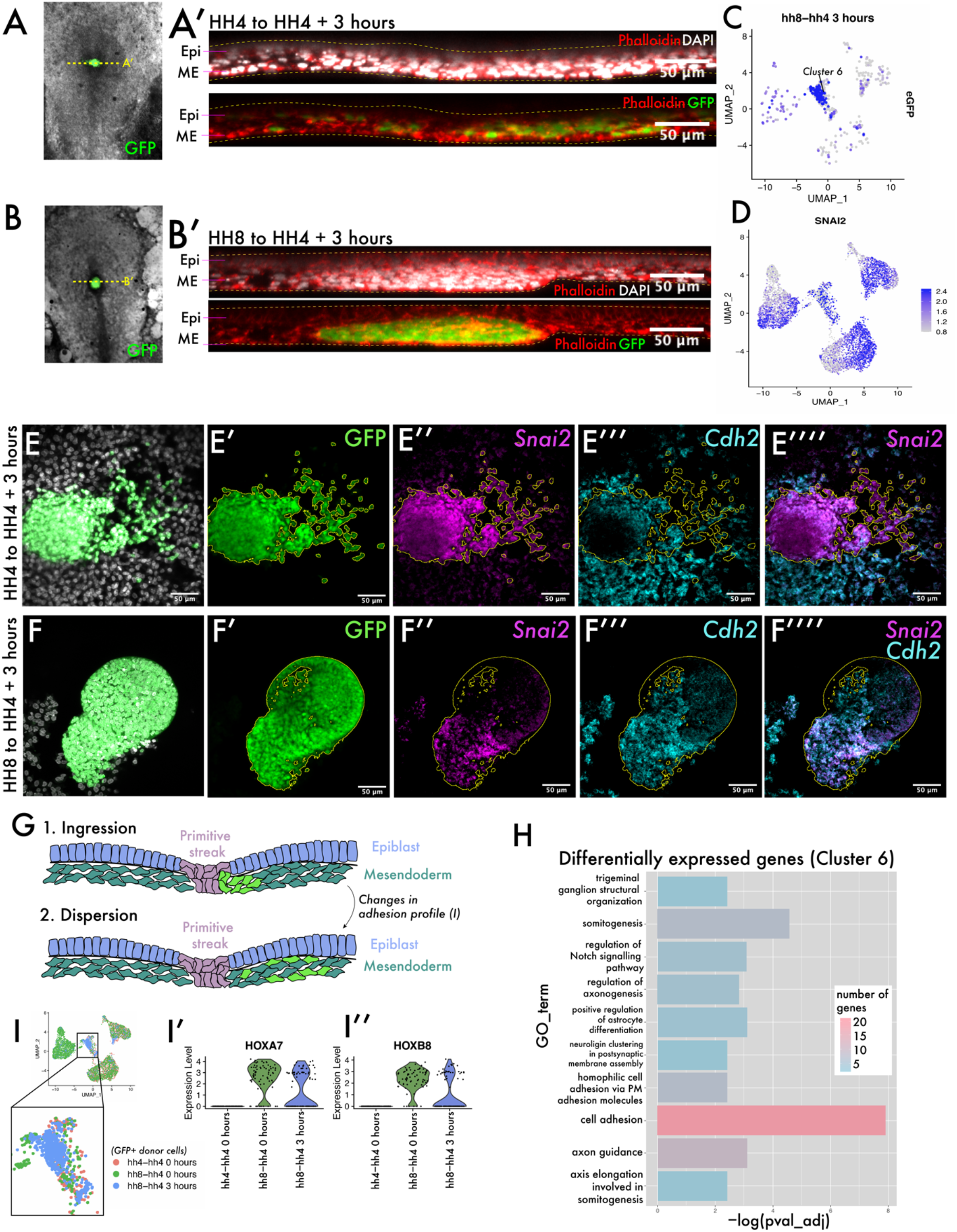
The transition from ingressed mesoderm to individual dispersed cells is an important step in gastrulation, associated with changes in cell adhesion. (A) and (B) are composite GFP and brightfield images of representative (HH4-HH4 and HH8-HH4 respectively) grafts at 3 hours after grafting. The dotted yellow lines in (A) and (B) represent the plane of section shown in (A’) and (B’) respectively. (A’) and (B’) are summed slice projections over ∼10μm of transverse re-sliced Z-stacks of the grafts in (A) and (B) respectively, fixed and stained with phalloidin (red) and DAPI (white). Labels show the position of the epiblast (‘epi’) and mesendodermal (‘ME’) cell layers in the axis. Images in (A’) and (B’) are oriented with dorsal at the top and ventral at the bottom. In both cases, GFP-expressing cells are found in the ventral ME cell layer. (C) UMAP plot for HH8-HH4 scRNA-seq sample with expression of *eGFP* shown, demonstrating the localisation of these cells in cluster 6 of the dataset. (D) UMAP plot of the full combined dataset showing *Snail2 (Snai2)* expression. Note the expression of *Snai2* in the central clusters of the plot. (E) and (F) are summed slice Z-projections of HH4-HH4 and HH8-HH4 grafts stained by multiplexed HCR for the EMT markers *Snai2* and *N-Cadherin (Cdh2*). (E) and (F) show DAPI, (E’) and (F’) show GFP fluorescence (donor tissue), (E’’) and (F’’) show *Snai2* expression, (E’’’) and (F’’’) show *Cdh2* expression and (E’’’’) and (F’’’’) show *Snai2* and *Cdh2* expression. The yellow outlines in panels (E) and (F) outline the location of GFP donor positive tissue. (G) Schematic showing the transition in mesoderm development from ingressed mesodermal progenitors to dispersed mesodermal progenitors in the ventral mesendodermal compartment, with the grafted green cells interspersed with surrounding unlabelled host tissue. (H) Gene Ontology (GO) term analysis of genes differentially expressed in cluster 6 of the dataset (the cluster in which the majority of HH8 grafted cellqs accumulate at 3 hours after grafting to the HH4 environment), represented here as a bar chart where the negative log of the adjusted p-value for each GO term is plotted. The colour of the bar represents the number of genes within the list of cluster 6 differentially expressed genes that are annotated with that GO term. (I) Schematic showing the cells utilised to look for differential expression between HH8 and HH4-derived cells in the central populations (clusters 6, 8 and 12). (I’) and (I’’) are violin plots showing the expression of the genes *HoxA7* and *HoxB8* in these cells by sample (note that only donor GFP+ cells are included in this plot). *Scale bars in A’, B’, E-F represent 50μm*.

To investigate what characterises the cell state that HH8 cells are stalled in, we performed Gene Ontology (GO) analysis on the list of genes differentially expressed in cluster 6 that are present across samples. Cluster 6 is where the majority of HH8 cells are found at 3 hours after grafting. We found a clear enrichment of genes associated with cell adhesion in this cluster (*Figure 3H*, full gene lists in *Supplementary Table 2*). To ask if these genes are differentially expressed between HH4 and HH8-derived cells (and thus could account for the different behaviour of these two populations upon grafting) we subsetted the scRNA-seq dataset to clusters 6, 8 and 12 only, annotated cells as either HH8 or HH4-derived (based on *eGFP* expression) and looked for genes differentially expressed between the two groups (schematic in *Figure 3I*). Note that there are differences between cell cycle state of HH8 and HH4 cells (*Supplementary Figure 2B, D-E)* and so we removed cell cycle-associated genes from the analysis. We found no significant difference in the expression of genes annotated with the ‘Cell Adhesion’ GO term identified in *Supp. Table 2* (*Supplementary Figure 2A*), suggesting that different expression of adhesion-related gene is not able to account for the different behaviour of HH8 and HH4 populations upon grafting. The most significantly enriched GO term in this analysis is ‘anteroposterior pattern specification’ (*Supplementary Figure 2C, Supplementary Table 3*), which includes the *Hox* genes *Hoxa7* and *Hoxb8*. These genes exhibit substantial expression in HH8-derived cells but no expression in HH4-derived cells (*Figure 3I’, I’’*). Together, the results from our scRNA-seq experiment show that upon grafting, HH8 cells initially occupy a region of gene expression space that is distinct from HH4 cells, but within three hours undergo transcriptomic changes that place them in a state similar to surrounding host tissue, with the exception of *Hox* gene expression differences.

### Both HH4 and HH8 cells migrate in an ex vivo culture system but with different dynamics

Having identified that HH8 grafts are delayed in their axis contribution at the level of cell migration from the graft site, we wondered whether this is an outcome of interaction between the donor and host tissue in this context or simply intrinsic to the grafted tissue. To test this, we developed an *ex vivo* culture system where MSP tissue was dissected and cultured on fibronectin-coated glass dishes and found that cells readily migrate as a sheet (*Supplementary Figure 3*). Both HH4 and HH8 MSP tissue migrates on fibronectin (*Figure 4A, C)*, so we used Particle Image Velocimetry (PIV) to characterise the dynamics of this process in more depth. Representative explant velocity fields are shown in *Figure 4B* and *4D* for HH4 and HH8 explants respectively. HH4 explants undergo a rapid expansion in area with no or little lag time before this expansion begins (*Figure 4E*) whereas HH8 tissue does not show a substantial increase in area for the first 10-15 hours of imaging, upon which time it expands relatively slowly (*Figure 4E*). The same trends can be seen in the velocity data, with HH4 explants showing an initial average velocity of 0.2*μ*m min^−1^and a maximal average velocity of ∼0.27*μ*m min^−1^at 9-10 hours of culture (*Figure 4F*). In contrast, HH8 tissue has a relatively constant velocity of ∼0.04*μ*m min^−1^rising to ∼0.1*μ*m min^−1^at 14-15 hours of culture (*Figure 4F*). Thus, HH8 tissue shows a slower increase in area (and corresponding lower velocity of tissue flow) than HH4 tissue, as well as a substantial lag time before spreading. This suggests that HH8 tissue intrinsically has a weaker proclivity to migrate than HH4 tissue and importantly can account for the lack of early migration we observe in heterochronic HH8 to HH4 grafts.

**Figure 4:**
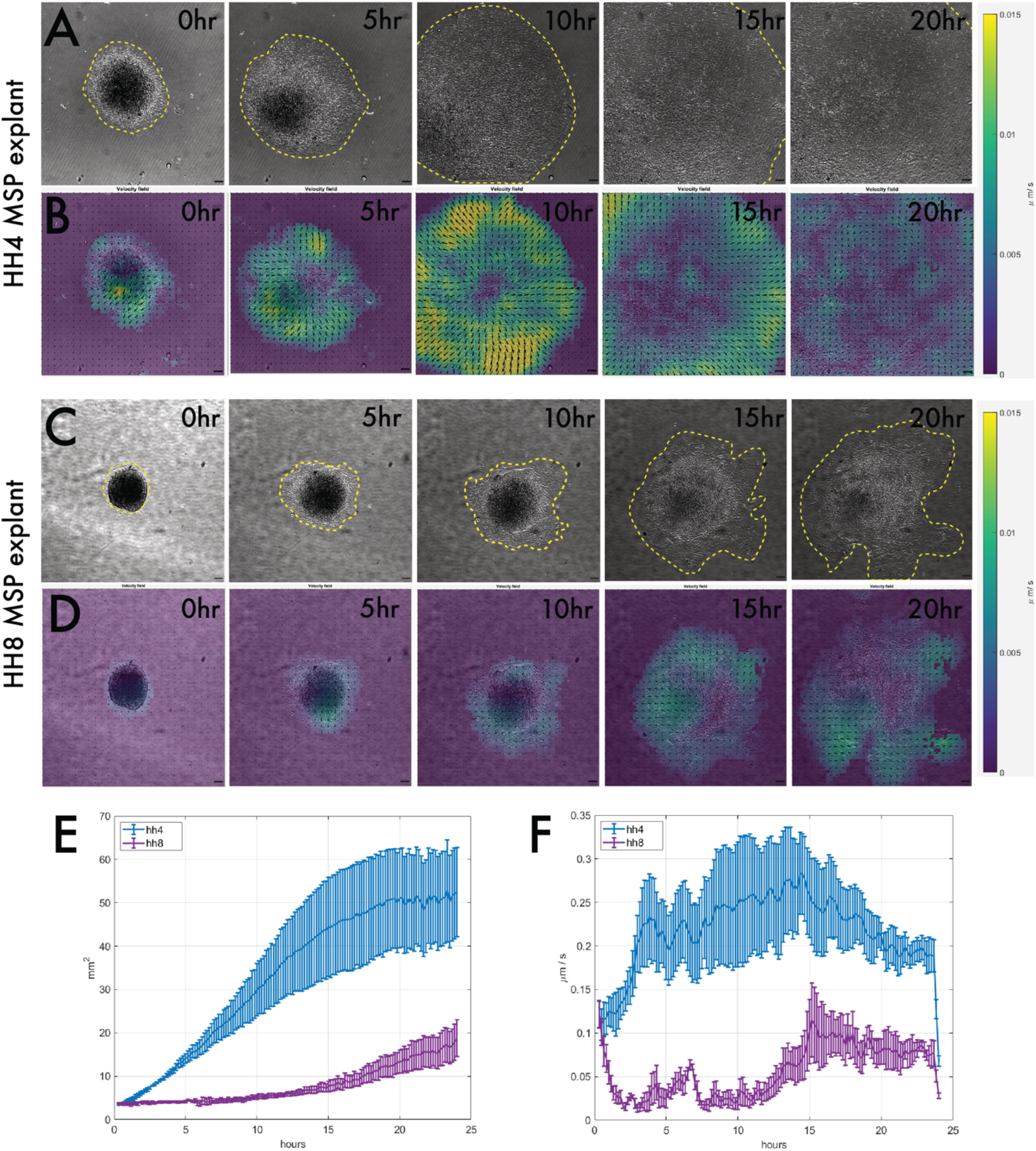
Explanting MSP tissue on fibronectin reveals that HH4 and HH8 MSP tissue intrinsically migrate with different dynamics. (A) and (C) are brightfield stills taken from timelapse movies of MSP tissue migrating on fibronectin, with (A) showing a HH4 explant and (C) showing a HH8 explant. Yellow dotted outlines delineate the edge of the migrating explant sheet. (B) and (D) are velocity field stills for the corresponding images in (A) and (C) respectively, calculated using PIV. The colour of each interrogation window denotes the velocity of movement (scale on right) between the current time step and the previous one, and the directionality of each vector arrow similarly represents the direction of tissue flow. (E) Plot showing the mean area for several explants of each class (n=5 for HH4 and n= 3 for HH8). (F) Plot showing the mean velocity at each timepoint for several explants of each class (n=5 for HH4 and n= 3 for HH8*). In (E) and (F), error bars represent the standard error. Scale bars in (A)-(D) represent 50μm*.

### Hox gene expression is intrinsically regulated in both grafts and explants

To further explore the intrinsic capability of MSP explants to regulate their developmental timing, we next aim to elucidate whether *in vivo Hox* gene expression is also tissue intrinsic. We identified a larger group of *Hox* genes with differential expression in the MSP region at HH4 and HH8 by screening by RT-qPCR and HCR staining (*Supplementary Figure 4*). These genes include *Hoxa2, Hoxa3* and *Hoxa6*, all of which are not expressed in the HH4 MSP region but expressed in the HH8 MSP region. We performed HCR for *Hoxa2, Hoxa3* and *Hoxa6* at 3, 6, 12 and 24 hours after grafting to ask whether HH8 tissue maintains expression of these genes upon grafting or downregulates their expression to match the surrounding host environment. We found that invariably, HH8 MSP cells retain the donor *Hox* profile upon grafting, suggesting that *Hox* gene expression may be regulated intrinsically at the population level (*Figure 5A-E*). As part of these experiments, we also performed grafts with variable sized pieces of donor tissue (*Supplementary Figure 5*). We found that at the smallest graft size (eighth size, approximately 6000μm^2^initial area), donor cells had spread substantially from the initial graft site (*Figure 5C)* whereas all larger graft sizes remained as a single cluster of cells (*Figure 5B*, example shown here is a quarter sized graft, approximately 12500μm^2^initial area). Interestingly, in the smallest grafts, individual GFP+ cells could be seen intermixed with wild-type host (GFP) tissue (*Figure 5C*, enlarged in *ii-iv)*. Notably, even these isolated HH8 cells still maintained expression of the HH8 *Hox* genes *Hoxa3* and *Hoxa6* (*Figure 5Cii-iv*), further supporting the intrinsic regulation of *Hox* gene expression in these cells. Additionally, this result also suggests that *Hox* gene expression can be uncoupled from cell dispersion, suggesting that *Hox* gene expression is not the only determinant of the timing of cell allocation to the body axis.

**Figure 5:**
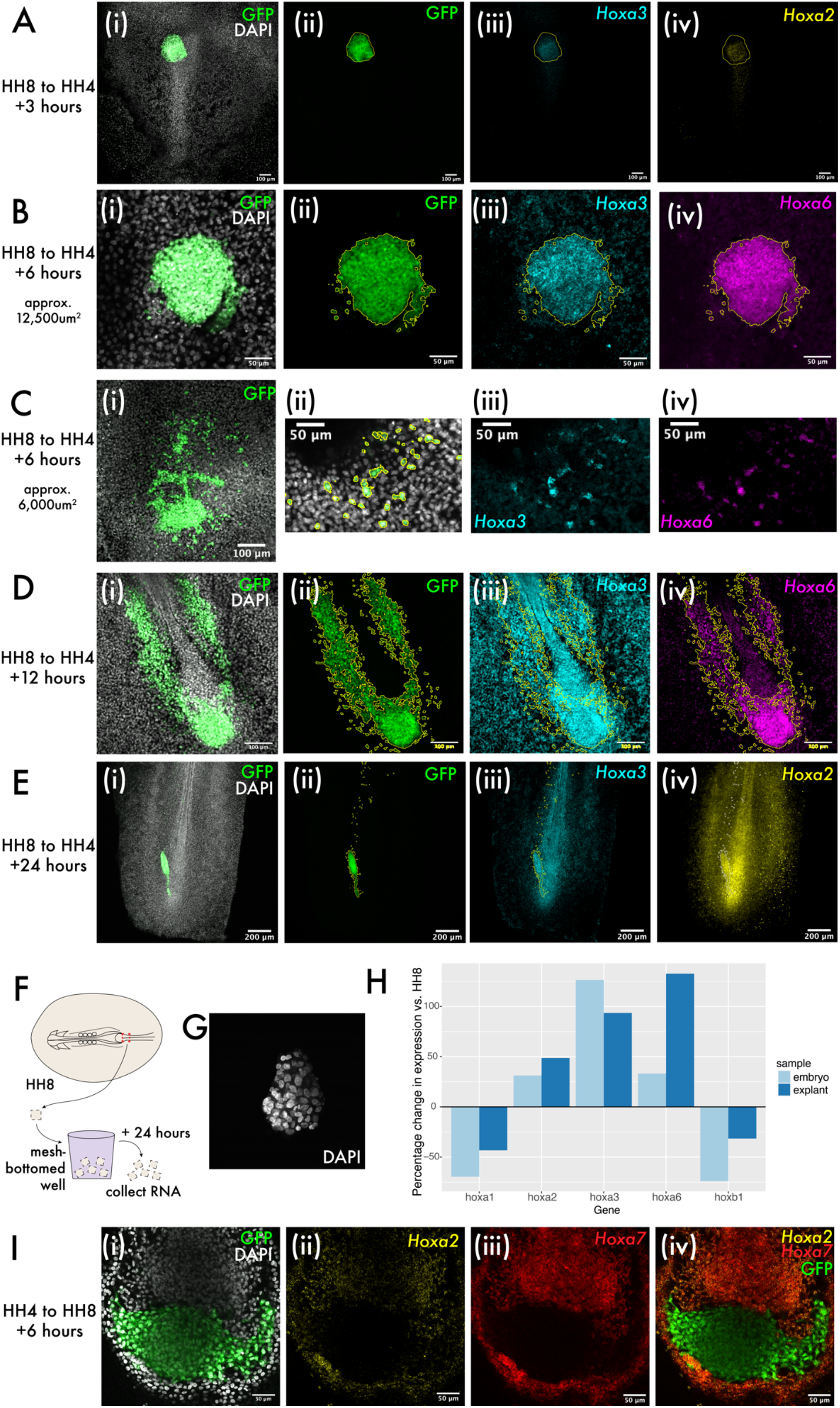
Hox expression is intrinsically regulated in both grafts and explants but can be uncoupled from cell dispersion. (A)-(E) show heterochronic HH8 to HH4 grafts at 3-, 6-, 6-, 12- and 24-hours post-grafting respectively, with each image showing the cells in the progenitor population (tailbud in (D) and (E)). Panel (i) of (A)-(E) shows composite DAPI and GFP summed slice projections. (ii), (iii) and (iv) are the same images as single channel summed slice projections, showing the location of the donor GFP+ cells, *Hoxa3* mRNA and *Hoxa2* or *Hoxa6* mRNA expression respectively (please see labels in panel for gene). In (C), (ii)-(iv) are enlarged images from a region within (C). In (A)-(E), yellow or white lines outline the location of GFP+ (donor) tissue. (F) Schematic showing explant experiment set-up. (G) DAPI stained HH8 explant after 24 hours of culture shown here as a summed slice projection. (H) Bar chart showing expression of five different *Hox* genes in embryonic MSP regions and HH8 explants after 24 hours, represented relative to the endogenous expression in MSP regions at HH8. (I) HH4 to HH8 graft at 6 hours after grafting, stained for *Hoxa2* and *Hoxa7* expression. (i) shows GFP and DAPI fluorescence, (ii) shows *Hoxa2* expression, (iii) shows *Hoxa7* expression and (iv) shows *Hoxa2, Hoxa7* and GFP. *Scale bars in (E) represent 200μm, in (A), (Ci) and (D) represent 100μm, and in (B), (Cii-iv) & (I) represent 50μm. All fluorescence images are presented as summed slice Z-*projections.

To investigate the intrinsic regulation of *Hox* expression further, we used explant culture to ascertain whether there is a role for the embryonic environment in progression of *Hox* expression. Explants were made of the HH8 MSP region and cultured for 24 hours as floating explants *(Figure 5F*) before RNA extraction. At 24 hours of culture, explants were maintained as cohesive floating pieces of tissue (*Figure 5G*). RT-qPCR for various *Hox* genes was performed to test for differences between *Hox* expression at 0 and 24 hours after explanting. In each case, this difference was compared to the endogenous change in expression in the MSP region over an equivalent period of time – 24 hours after HH8 embryos are HH14, so MSP tissue at HH14 was used to represent the endogenous change in expression (*Figure 5H*). Consistent with a progression of the Hox timer in the cells of the explants over 24 hours, we observed an increase or decrease in *Hox* expression consistent with the normal dynamic for each gene surveyed (*Figure 5H*). This data provides support for population-intrinsic regulation of Hox gene expression in MSPs, with this timer able to progress outside of the embryonic environment.

Finally, we performed HH4 to HH8 grafts to ascertain whether there was any asymmetry in the extrinsic regulation of *Hox* gene expression depending on whether tissue was grafted from older to younger or younger to older embryos. We found that in this instance – where HH4 donor tissue does not express *Hoxa2* or *Hoxa7 in vivo*, but host HH8 tissue does – there was no evidence for the upregulation of these genes in donor tissue at 6 hours after grafting (*Figure 5I*), further supporting an intrinsic mode of regulation of *Hox* gene expression in this context. Together, the experiments described here suggest that *Hox* gene expression in the MSP population is intrinsically regulated and not subject to reprogramming to match the surrounding host environment.

## Discussion

### Mesodermal dispersion is a key step in avian gastrulation

In this work, we first sought to understand at what point in the developmental process of progenitor contribution to the body axis heterochronic grafts of cells from older to younger embryos are delayed. The delayed contribution of axial progenitor cells to the axis that we observe is consistent with previous studies, including mouse tailbud grafts (Tam and Tan, 1992) and chicken primitive streak grafts (Iimura and Pourquie, 2006). Based on single cell RNA-sequencing and *in vivo* gene expression analyses, we propose that MSP contribution to the body axis can be conceptualised as consisting of three successive phases: ingression, dispersion, and migration (*Figure 3G*). In the first step, mesodermal progenitor cells of the primitive streak undergo a classical epithelial-to-mesenchymal transition (EMT) and ingress from the dorsal cell layer to the ventral one. This EMT involves a well-characterised set of molecular players including the upregulation of *Snail2* (*Slug*) and a switch from E-Cadherin (*Cdh1*) to N-Cadherin (*Cdh2*) expression (Cano et al., 2000; Dady et al., 2012; Nieto et al., 1994; Ohkubo and Ozawa, 2004). Once cells enter the mesendodermal cell layer, substantial and rapid cell dispersion occurs in both the anteroposterior and mediolateral axes. Finally, cells must move to the final site of somite formation (Bénazéraf et al., 2010; Yang et al., 2002). It is prior to the second stage in this process, dispersion, at which heterochronically grafted HH8 cells are paused, resulting in the population being maintained as a distinct cluster of cells in the ME and not dispersing readily as control grafts do. These processes are recapitulated in the explant culture system we describe, where both HH4 and HH8 MSP populations ultimately migrate on fibronectin but HH8 populations do so with a delay and at lower velocities. Thus, delayed dispersion of HH8 tissue in heterochronic grafts to HH4 embryos can be accounted for by intrinsic migratory properties of the HH8 tissue.

As discussed in the introduction, the choice of HH4 and HH8 tissue as the focus for this project is interesting in the context of primary versus secondary gastrulation: as mentioned, the occipital somites form simultaneously from tissue that ingresses during primary gastrulation, whereas trunk somites form periodically through the progressive segmentation of a presomitic mesoderm (PSM). Here, we have shown that HH4 MSP tissue (pre-node regression) and HH8 MSP tissue (post-node regression) exhibit substantially different migratory dynamics, despite forming a continuous progenitor population. This concept of evolving cell behaviour and properties over time is consistent with the transcriptional changes that have been described in axial progenitor regions (Wymeersch et al., 2019).

### Hox gene regulation is population intrinsic for somite progenitors

Vertebrate embryos express Hox genes with a special dynamic – each paralogous cluster of Hox genes is expressed with spatiotemporal collinearity, meaning that they are transcribed in a strict 3’ to 5’ order both in space and time. It is known that the location within the cluster defines the timing of expression of each gene, that *Hox* clusters undergo epigenetic remodelling to increase DNA accessibility in a 3’ to 5’ direction, and that the initial induction of Hox gene expression at gastrulation is dependent upon Wnt signalling (Deschamps and Duboule, 2017; Izpisúa-Belmonte et al., 1991; Kmita and Duboule, 2003; Liu et al., 1999; Moreau et al., 2019; Neijts et al., 2016; Tschopp et al., 2009). We find evidence for a population-intrinsic mechanism of *Hox* gene expression – once HH8 cells are expressing a given *Hox* gene, they will maintain this expression profile in a novel HH4 environment. The maintenance of Hox gene expression in a new environment (a younger embryo) is consistent with a result reported by Iimura and Pourquie, 2006, wherein *Hoxb9* expression was shown to persist at 6 hours after heterochronic grafting of streak cells. However, our results differ from those reported by McGrew et al., 2008, who observed reprogramming of *Hox* gene expression in HH15 tailbud chordoneural hinge cells transplanted to the equivalent region of the HH8 embryo – grafted cells no longer express the *Hox* gene *Hoxa10* at eight hours after grafting. This difference may reflect the different cell populations studied or variation in the contributions of extrinsic and intrinsic influences to Hox regulation over the period of primary body axis formation; it is possible that there is a transition from intrinsic to a more extrinsic mode of *Hox* regulation later in primary body axis development. Alternatively, it may be explained by the relative mismatch between donor and host tissue. In the experiments of McGrew et al. (2008), *Hoxa10* expression is absent from the progenitor region at HH8-HH10 but expressed at HH15. *Hoxa10* expression is first detected in the progenitor region at HH12 (17 somite stage) (Denans et al., 2015), corresponding to a theoretical 20-hour lag time before expression of *Hoxa10* in host tissue of HH8 to HH15 grafts. Conversely, our HH8 to HH4 grafts experience a much shorter lag period before host expression of the *Hox* genes surveyed – expression of *Hoxa2* for example is evident in host tissue within 6 hours. Additional grafts and additional *Hox* genes must be studied to ascertain which of these possible explanations is most accurate.

We have seen that the difference in behaviour between homochronic and heterochronic grafts correlates with the persistence of the HH8 *Hox* gene expression profile, raising the possibility that the HH8 *Hox* profile may influence the transition from mesenchyme to invasive mesenchyme. These results differ somewhat from observations after over-expressing *Hox* genes in primitive streak tissue, where cells were found to be delayed at the point of ingression through the streak (Iimura and Pourquie, 2006). In this study, cells over-expressing more 5’ Hox genes remained in the epiblast or primitive streak for longer before entering the ventral mesendoderm cell layer. This difference may derive from the use of physiological levels of *Hox* transcript in our study versus overexpression of transcript in the over-expression experiments described.

### Population size-related inputs on cell allocation to the axis

Our work shows that smaller grafts of older progenitor populations to primitive streak stage embryos do not remain as a single mass but instead show substantial mixing with host tissue, like homochronic grafts. The observation of graft size influencing outcome resembles previous findings that grafts of neural tissue have different outcomes dependent upon graft size (Couly et al., 1998; Guthrie et al., 1992; Trainor and Krumlauf, 2000). A common differential outcome is the maintenance of donor gene expression in larger grafts but not in instances when donor cells are isolated or in smaller groups (Schilling et al., 2001; Trainor and Krumlauf, 2000). Interestingly, in our experiments, we observe maintenance of the donor *Hox* profile even when cells are not contacting any other donor cells, suggesting that *Hox* gene expression in this context may be truly cell intrinsic. Cell intrinsic timers have been described in diverse contexts when individual cells are isolated from their developmental contexts, including the somitogenesis clock (Webb et al., 2016), the *Drosophila* neuroblast clock (Grosskortenhaus et al., 2005) and oligodendrocyte precursor cell (OPC) differentiation timing (Temple and Raff, 1986).

Together, our work has shown that heterochronic somitic progenitor grafts from older to younger embryos are delayed at a previously uncharacterised stage that we term dispersion, remaining as a single group of cells rather than mixing with surrounding host tissue. This difference occurs after ingression through the primitive streak and correlates with the intrinsic regulation of *Hox* gene expression, such that expression is robustly maintained upon grafting to the HH4 environment. We find that despite this maintenance, HH8 cells can disperse if they are transplanted in relatively small groups, suggesting that *Hox* gene expression is not the only input timing cell contribution to the primary body axis. Explant culture has shown that the delayed dispersion of larger grafts can be accounted for by intrinsic migratory dynamics of HH8 tissue. Overall, our results are consistent with an emerging view of vertebrate developmental timing where inputs across length scales as well as a combination of intrinsic and extrinsic timer mechanisms give rise to a stereotyped order and timing of events in development (Busby and Steventon 2021; Chinnaiya et al., 2014; Matsuda et al., 2020; Rayon et al., 2020; Pickering et al., 2018; Saiz-Lopez et al., 2015, 2017).

## Acknowledgements

L.B. was supported by a BBSRC DTP Studentship, G.S.N. is supported by a research grant from the Leverhulme Trust (RG93881), and B.S. was supported by a Henry Dale Fellowship jointly funded by the Wellcome Trust and the Royal Society (109408/Z/15/Z). We thank Alexandra Neaverson, Val Wilson, and Kate McDole for valuable feedback.

## Author contributions

Conceptualization, L.B. and B.S.; Methodology, L.B. and B.S.; Investigation, L.B. and G.S.N.; Writing – Original Draft, L.B. and B.S.; Visualization, L.B. and G.S.N.; Funding Acquisition, L.B., B.S, G.S.N.

## Data availability

The single cell RNA-sequencing dataset generated in this work is available via the NCBI Gene Expression Omnibus (GEO) under the accession code GSE224169.

## Materials & Methods

### Chicken husbandry

Bovans brown chicken eggs (Henry Stewart & Co., Norfolk) and GFP transgenic chicken eggs (Roslin Institute, Edinburgh) were stored upon arrival at 18 degrees Celsius until use, then incubated at 37 degrees Celsius in a humidified incubator to obtain embryos of an appropriate stage according to Hamburger & Hamilton, 1951.

#### Tissue fixation

Embryos were fixed for analysis by dissection in PBS (Sigma-Aldrich) and fixation in 4% PFA (Sigma-Aldrich) in PBST either overnight at 4 degrees Celsius or for 1 hour at room temperature (RT).

### Experimental embryology

#### Grafting

Homochronic and heterochronic grafts were performed with host embryos in New Culture (New, 1955). Donor GFP-expressing tissue pieces (McGrew *et al*., 2008) containing ∼1000-1200 cells were dissected from the embryo in PBS, then transferred to the host embryo in new culture with a glass capillary tube. The graft was carefully positioned at the graft site under PBS (Sigma-Aldrich) immersion using an eyelash tool, and excess PBS removed, before re-incubation at 37 degrees Celsius.

#### Floating explant culture

Explants of the HH8 medial somite progenitor region were prepared as follows. Wild-type embryos were removed from the egg, washed in PBS, and the MSP region isolated using a micro-dissecting knife. Explants were cultured in Medium 199 (Gibco) in Millicell cell culture inserts (based on methods in Streit and Stern, 1999) for 24 hours, before collection for either fixation or RNA extraction.

#### Fibronectin (adhesive) explant culture

Glass-bottomed imaging dishes (Mattek) were coated with human fibronectin protein (1mg/mL stock solution diluted 1:40 in PBS) by pipetting the solution onto the dish and incubation at 37ºC for 1-3 hours. Fibronectin solution was removed, and the dish allowed to dry for 15 minutes at 37ºC. Explants of the HH4 or HH8 MSP region were prepared as follows. Wild-type embryos were removed from the egg, washed in PBS, and the MSP region isolated using a micro-dissecting knife. Explants were placed on the fibronectin coated dish and oriented mesendoderm (ventral) side down using an eyelash tool and cultured in GMEM (Gibco) + 10% FBS + 1% Pen/Strep.

### Molecular biology

#### Hybridisation Chain Reaction (HCR)

RNA expression was visualised using third-generation HCR (Choi et al., 2018). Embryos were rehydrated through a descending methanol series, washed twice in PBST, then digested in Proteinase K (10μg/mL) for 5-10 minutes at room temperature. Embryos were then incubated with 4% PFA in PBST for 20 minutes, washed twice in 5X SSCT for 10 minutes each, and then incubated in HCR hybridisation buffer (Molecular Instruments) for 30 minutes at 37ºC. HCR probes (Molecular Instruments) were added to hybridisation buffer at 4ρmol/mL and embryos were allowed to incubate in this probe solution overnight at 37ºC. The next day, embryos were washed four times for 15 minutes in HCR wash buffer (Molecular Instruments) at 37ºC, then washed twice in 5X SSCT at room temperature for 10 minutes. Embryos were incubated in HCR amplification buffer (Molecular Instruments) for 5 minutes. Fluorescently labelled HCR hairpins (Molecular Instruments) were snap-cooled by incubation at 95ºC for 90 seconds before cooling to room temperature in darkness. HCR hairpins were added to HCR amplification buffer at 60ρmol/mL each, then embryos were incubated in this hairpin solution overnight at room temperature in darkness. Hairpin solution was removed, and embryos were washed in 5X SSCT for 10 minutes three times at room temperature. DAPI staining was performed by incubation of embryos in DAPI (1:1000) in 5X SSCT for 30 minutes at room temperature. Finally, DAPI solution was washed off by three 10-minute washes in 5X SSCT at room temperature.

#### Phalloidin staining

Fixed embryos were stained for F-actin using Alexa 568 conjugated phallodin (Thermo-Fisher). Embryos were incubated in a 1:1000 dilution of phallodin in PBS for 1-3 hours at room temperature before washing in PBS.

#### RNA extractions

Tissue was dissected in cold PBS and transferred to 1mL cold Trizol (Invitrogen), homogenised using a pestle, and allowed to incubate at room temperature for 5 minutes. 200μL chloroform was added and the mixture was incubated for 3 minutes at room temperature. Samples were centrifuged for 15 minutes at 4ºC, and the upper aqueous phase was retained and transferred to a new tube. 500μL isopropanol was added to the aqueous phase, incubated for 10 minutes at RT, and then centrifuged for 10 minutes at 4ºC. The supernatant was discarded, and the pellet washed with 75% ethanol, before resuspension in RNase-free water and incubation at 55ºC for 15 minutes.

#### RT-qPCR

RNA was reverse transcribed to cDNA using SuperScript III reverse transcriptase at 50ºC for 60 minutes. RNA was removed by RNase H digestion for 20 minutes at 37ºC. QPCR was performed using the Quiagen Rotor-Gene Q machine, with SYBR Green Mastermix. Reactions were run in triplicate. Gene expression was quantified by standard curve analyses using serial dilution series of HH7 whole embryo cDNA. The threshold cycle (Ct) was calculated for each dilution in the series (using a threshold of 0.02) and used to produce a log dilution-Ct graph for each gene, allowing calculation of the relative concentration of transcript in each unknown sample. Primer sequences are available in *Supplementary Table 4*.

### Single cell RNA-sequencing

Samples were prepared for single-cell RNA sequencing as follows: tissue was dissected in cold PBS and washed thrice in cold PBS. PBS was replaced with dissociation solution (Trypsin + 0.05% EDTA, Gibco) and samples were incubated at 37ºC for 15 minutes, with trituration every 5 minutes. The dissociation reaction was terminated by addition of 10% Foetal Bovine Serum (FBS). Samples were spun down at 4ºC for 5 minutes and the supernatant discarded. Each cell pellet was resuspended in PBS + 0.25% BSA and passed through a 40μm pluriStrainer cell strainer (pluriSelect). cDNA libraries were prepared using the 10X Genomics 3’mRNA-seq workflow (10X Genomics) and sequenced using a NovaSeq instrument (Illumina). Data was processed using CellRanger (10X Genomics) to align reads to a reference transcriptome (produced using the galGal6 genome assembly published by Genome Reference Consortium) and produce matrices of gene expression. Matrices were then passed into Seurat in R (Satija et al., 2015) to perform clustering analyses and identify differential gene expression amongst clusters. Gene Ontology analysis was performed with the GeneCodis webtool (Carmona-Saez et al., 2007).

### Imaging

Live embryos were imaged on a fluorescent dissecting scope (Leica) in brightfield and GFP channels. Images were overlaid using GNU Image Manipulation Programme (GIMP) to produce the composite images presented in the paper. Confocal imaging of fixed and stained embryos was performed using a Zeiss LSM 700. For confocal imaging, embryos were mounted in Vectashield between two coverslips separated by double sided tape and sealed using clear nail polish. For the live imaging of adherent explants, a Nikon widefield scope was used with a heated chamber (37ºC).

### Particle Image Velocimetry (PIV)

The velocity fields at the surface of the explants were computed by digital PIV using PIVLab vs2.56 for MATLAB with two passes of 64 × 64 pixels and 32 × 32 pixels with a 50% overlap. To reduce the noise and remove small fluctuations, each interrogation window was averaged with its 7 by 7 neighbourhood, and with the same location in the previous and following time point. Velocity vectors outside of the explant were eliminated using a mask generated using Trainable Weka Segmentation, a Fiji plugin that allows training of machine learning models for per-pixel classification (Schindelin et al., 2012). The pixel size for this data is 1.29 μm and the time step is 10 minutes.

## References

1. Barak, H., Preger-Ben Noon, E., and Reshef, R. (2012). Comparative spatiotemporal analysis of Hox gene expression in early stages of intermediate mesoderm formation. Dev. Dyn. 241, 1637–1649.

2. Bell, G.W., Yatskievych, T.A., and Antin, P.B. (2004) GEISHA, a high throughput whole mount in situ hybridization screen in chick embryos. Devel. Dynamics 229: 677–687.

3. Bénazéraf, B., Francois, P., Baker, R.E., Denans, N., Little, C.D., and Pourquie, O. (2010). A random cell motility gradient downstream of FGF controls elongation of an amniote embryo. Nature 466, 1–5.

4. Brown, J., and Storey, K. (2000). A region of the vertebrate neural plate in which neighbouring cells can adopt neural or epidermal fates. Current Biology 10(14), 869–872.

5. Busby, L., and Steventon, B. (2021). Tissue tectonics and the multi-scale regulation of developmental timing. R. Soc. Interface Focus 11.

6. Busby, L., Saunders, D., Serrano Najera, G., and Steventon, B. (2022). Quantitative Experimental Embryology: A Modern Classical Approach. Journal of Developmental Biology 10(44).

7. Cano, A., Pérez-Moreno, M.A., Rodrigo, I., Locascio, A., Blanco, M.J., Del Barrio, M.G., Portillo, F., and Nieto, M.A. (2000). The transcription factor Snail controls epithelial-mesenchymal transitions by repressing E-cadherin expression. Nat. Cell Biol. 2, 76–83.

8. Carmona-Saez, P., Chagoyen, M., Tirado, F., Carazo, J.M., and Pascual-Montano, A. (2007). GENECODIS: a web-based tool for finding significant concurrent annotations in gene lists. Genome Biology 8:R3.

9. Catala, M., Teillet, M-A., Dr Robertis, E. M., and Le Douarin, N. M. (1996). A spinal cord fate map in the avian embryo: while regressing, Hensen’s node lays down the notochord and floor plate thus joining the spinal cord lateral walls. Development 122, 2599–2610.

10. Chinnaiya, K., Tickle, C., and Towers, M. (2014). Sonic hedgehog-expressing cells in the developing limb measure time by an intrinsic cell cycle clock. Nat. Commun. 5, 1–8.

11. Choi, H.M.T., Schwarzkopf, M., Fornace, M.E., Acharya, A., Artavanis, G., Stegmaier, J., Cunha, A., and Pierce, N.A. (2018). Third-generation in situ hybridization chain reaction: multiplexed, quantitative, sensitive, versatile, robust. Development 145, dev165753.

12. Couly, G., Coltey, P. M., and Le Douarin, N. M. (1993). The triple origin of skull in higher vertebrates: a study in quail-chick chimeras. Development 117(2), 409–429.

13. Couly, G., Grapin-Botton, A., Coltey, P., Ruhin, B., and Le Douarin, N.M. (1998). Determination of the identity of the derivatives of the cephalic neural crest: incompatibility between Hox gene expression and lower jaw development. Development 125, 3445–3459.

14. Dady, A., Blavet, C., and Duband, J.L. (2012). Timing and kinetics of E-to N-cadherin switch during neurulation in the avian embryo. Dev. Dyn. 241, 1333–1349.

15. Darnell, D.K., Kaur, S., Stanislaw, S., Davey, S., Konieczka, J.H., Yatskievych, T.A., and Antin, P.B. (2007). GEISHA: An In situ hybridization gene expression resource for the chicken embryo. Cytogenet. Genome Res. 117, 30–35.

16. Denans, N., Iimura, T., City, K., States, U., City, K., States, U., Biology, C., City, K., States, U., States, U., et al. (2015). Hox genes control vertebrate body elongation by collinear Wnt repression. Elife 4:e04379, 1–33.

17. Deschamps, J., and Duboule, D. (2017). Embryonic timing, axial stem cells, chromatin dynamics, and the Hox clock. Genes Dev. 31, 1406–1416.

18. Dias, A. S., De Almeida, I., Belmonte, J. M., Glazier, J. A., and Stern, C. D. (2014). Somites without a clock. Science 343(6172), 791–795.

19. Duboule, D. (2022). The (unusual) heuristic value of Hox gene clusters; a matter of time? Dev. Biol. 484, 75–87.

20. Ebisuya, M., and Briscoe, J. (2018). What does time mean in development? Development 145, dev164368.

21. Fulton, T., Speiss, K., Thomson, L., Wang, Y., Clark, B., Hwang, S., Paige, B., Verd, B., and Steventon, B. (2022). Cell Rearrangement Generates Pattern Emergence as a Function of Temporal Morphogen Exposure. Pre-print at BioRxiv, accessed 05.05.2023.

22. Gray, S. D., and Dale, J. K. (2010). Notch signalling regulates the contribution of progenitor cells from the chick Hensen’s node to the floor plate and notochord. Development 137, 561–568.

23. Grosskortenhaus, R., Pearson, B.J., Marusich, A., and Doe, C.Q. (2005). Regulation of temporal identity transitions in drosophila neuroblasts. Dev. Cell 8, 193–202.

24. Guthrie, S., Muchamore, I., Kuroiwa, A., Marshall, H., Krumlauf, R., and Lumsden, A. (1992). Neuroectodermal autonomy of Hox-2.9 expression revealed by rhombomere transpositions. Nature 356, 157–159.

25. Hamburger, V., and Hamilton, H.L. (1951). A series of normal stages in the development of the chick embryo. J. Morphol. 88, 49–92.

26. Huang, R., Zhi, Q., Patel, K., Wilting, J., and Christ, B. (2000). Contribution of single somites to the skeleton and muscles of the occipital and cervical regions in avian embryos. Anatomy and Embryology 202, 375–383.

27. Iimura, T., and Pourquie, O. (2006). Collinear activation of Hoxb genes during gastrulation is linked to mesoderm cell ingression. Nature 442, 1–4.

28. Iimura, T., Yang, X., Weijer, C.J., and Pourquié, O. (2007). Dual mode of paraxial mesoderm formation during chick gastrulation. Proc. Natl. Acad. Sci. U. S. A. 104, 2744–2749.

29. Izpisúa-Belmonte, J.C., Falkenstein, H., Dollé, P., Renucci, A., and Duboule, D. (1991). Murine genes related to the Drosophila AbdB homeotic genes are sequentially expressed during development of the posterior part of the body.EMBO J. 10, 2279–2289.

30. Joshi, P., Darr, A.J., and Skromne, I. (2019). CDX4 regulates the progression of neural maturation in the spinal cord. Dev. Biol. 449, 132–142.

31. Kmita, M., and Duboule, D. (2003). Organizing axes in time and space; 25 years of colinear tinkering. Science 301, 331–333.

32. Lim, T. M., Lunn, E. R., Keynes, R. J., and Stern, C. D. (1987). The differing effects of occipital and trunk somites on neural development in the chick embryo. Development 100(3), 525–533.

33. Liu, P., Wakamiya, M., Shea, M.J., Albrecht, U., Behringer, R.R., and Bradley, A. (1999). Requirement for Wnt3 in vertebrate axis formation. Nat. Genet. 22, 361–365.

34. Matsuda, M., Hayashi, H., Garcia-Ojalvo, J., Yoshioka-Kobayashi, K., Kageyama, R., Yamanaka, Y., Ikeya, M., Toguchida, J., Alev, C., and Ebisuya, M. (2020). Species-specific segmentation clock periods are due to differential biochemical reaction speeds. Science 369, 1450–1455.

35. McGrew, M.J., Sherman, A., Lillico, S.G., Ellard, F.M., Radcliffe, P.A., Gilhooley, H.J., Mitrophanous, K.A., Cambray, N., Wilson, V., and Sang, H. (2008). Localised axial progenitor cell populations in the avian tail bud are not committed to a posterior Hox identity. Development 135, 2289–2299.

36. Moreau, C., Caldarelli, P., Rocancourt, D., Roussel, J., Denans, N., Pourquie, O., Moreau, C., Caldarelli, P., Rocancourt, D., Roussel, J., et al. (2019). Timed Collinear Activation of Hox Genes during Gastrulation Controls the Avian Forelimb Position. Curr. Biol. 29, 35–50.

37. Neijts, R., Amin, S., Van Rooijen, C., Tan, S., Creyghton, M.P., De Laat, W., and Deschamps, J. (2016). Polarized regulatory landscape and Wnt responsiveness underlie Hox activation in embryos. Genes Dev. 30, 1937–1942.

38. Nieto, M.A., Sargent, M.G., Wilkinson, D.G., and Cooke, J. (1994). Control of cell behavior during vertebrate development by Slug, a zinc finger gene. Science 264, 835–839.

39. Ohkubo, T., and Ozawa, M. (2004). The transcription factor Snail downregulates the tight junction components independently of E-cadherin downregulation. J. Cell Sci. 117, 1675–1685.

40. Pickering, J., Rich, C.A., Stainton, H., Aceituno, C., Chinnaiya, K., Saiz-Lopez, P., Ros, M.A., and Towers, M. (2018). An intrinsic cell cycle timer terminates limb bud outgrowth. Elife 7, 1–15.

41. Psychoyos, D., and Stern, C. (1996). Fates and migratory routes of primitive streak cells in the chick embryo. Development 122(5), 1523–1534.

42. Raff, M. (2006). The mystery of intracellular developmental programmes and timers. Biochem. Soc. Trans. 34, 663–670.

43. Rayon, T., Stamataki, D., Perez-Carrasco, R., Garcia-Perez, L., Barrington, C., Melchionda, M., Exelby, K., Lazaro, J., Tybulewicz, V.L.J., Fisher, E.M.C., et al. (2020). Species-specific pace of development is associated with differences in protein stability. Science 369, eaba7667.

44. Rayon, T. (2023). Cell time: how cells control developmental timetables. Science Advances 9(10).

45. Rodrigues, S., Santos, J., and Palmeirim, I. (2006). Molecular characterization of the rostral-most somites in early somitic stages of the chick embryo. Gene Expression Patterns 6(7), 673–677.

46. Saiz-Lopez, P., Chinnaiya, K., Campa, V.M., Delgado, I., Ros, M.A., and Towers, M. (2015). An intrinsic timer specifies distal structures of the vertebrate limb. Nat. Commun. 6, 1–9.

47. Saiz-Lopez, P., Chinnaiya, K., Towers, M., and Ros, M.A. (2017). Intrinsic properties of limb bud cells can be differentially reset. Development 144, 479–486.

48. Satija, R., Farrell, J.A., Gennert, D., Schier, A.F., and Regev, A. (2015). Spatial reconstruction of single-cell gene expression data. Nat. Biotechnol. 33.

49. Schilling, T.F., Prince, V., and Ingham, P.W. (2001). Plasticity in zebrafish hox expression in the hindbrain and cranial neural crest. Dev. Biol. 231, 201–216.

50. Schindelin, J., Arganda-Carreras, I., Frise, E., Kaynig, V., Longair, M., Pietzsch, T., Preibisch, S., Rueden, C., Saalfeld, S., Schmid, B., et al. (2012). Fiji: An open-source platform for biological-image analysis. Nat. Methods 9, 676–682.

51. Schoenwolf, G. (1977). Tail (end) bud contributions to the posterior region of the chick embryo. Journal of Experimental Zoology 201(2), 227–245.

52. Selleck, M.A.J., and Stern, C.D. (1991). Fate mapping and cell lineage analysis of Hensen’s node in the chick embryo. Development 112, 615–626.

53. Streit, A., and Stern, C.D. (1999). Establishment and maintenance of the border of the neural plate in the chick: Involvement of FGF and BMP activity. Mech. Dev. 82, 51–66.

54. Tam, P.P.L., and Tan, S.S. (1992). The somitogenetic potential of cells in the primitive streak and the tail bud of the organogenesis-stage mouse embryo. Development 115, 703–715.

55. Temple, S., and Raff, M.C. (1986). Clonal analysis of oligodendrocyte development in culture: Evidence for a developmental clock that counts cell divisions. Cell 44, 773–779.

56. Trainor, P., and Krumlauf, R. (2000). Plasticity in mouse neural crest cells reveals a new patterning role for cranial mesoderm. Nat. Cell Biol. 2, 96–102.

57. Tschopp, P., Tarchini, B., Spitz, F., Zakany, J., and Duboule, D. (2009). Uncoupling time and space in the collinear regulation of Hox genes. PLoS Genet. 5, 1–12.

58. Webb, A.B., Lengyel, I.M., Jörg, D.J., Valentin, G., Jülicher, F., Morelli, L.G., and Oates, A.C. (2016). Persistence, period and precision of autonomous cellular oscillators from the zebrafish segmentation clock. Elife 5, 1–17.

59. Wymeersch, F. J., Skylaki, S., Huang, Y., Watson, J. A., Economou, C., Marek-Johnston, C., Tomlinson, S. R., and Wilson, V. (2019). Transcriptionally dynamic progenitor populations organised around a stable niche drive axial patterning. Development 146, dev168161.

60. Yang, X., Dormann, D., Munsterberg, A. E., and Weijer, C. J. (2002). Cell movement patterns during gastrulation in the chick are controlled by positive and negative chemotaxis mediated by FGF4 and FGF8. Developmental Cell 3, 425–437.

